# VE607 Stabilizes SARS-CoV-2 Spike In the “RBD-up” Conformation and Inhibits Viral Entry

**DOI:** 10.1101/2022.02.03.479007

**Authors:** Shilei Ding, Shang Yu Gong, Jonathan Grover, Mohammadjavad Mohammadi, Yaozong Chen, Dani Vézina, Guillaume Beaudoin-Bussières, Vijay Tailor Verma, Guillaume Goyette, Jonathan Richard, Derek Yang, Amos B. Smith, Marzena Pazgier, Marceline Côté, Cameron Abrams, Walther Mothes, Andrés Finzi, Christian Baron

**Affiliations:** Centre de recherche du CHUM, Montréal, QC, Canada; Department of Microbiology and Immunology, McGill University, Montreal, QC, Canada; Department of Microbial Pathogenesis, Yale University School of Medicine, New Haven, CT 06510, USA; Department of Biochemistry and Molecular Biology, Drexel University College of Medicine, Philadelphia, PA 19104, USA; Infectious Disease Division, Department of Medicine, Uniformed Services University of the Health Sciences, Bethesda, MD 20814-4712, USA; Département de Microbiologie, Infectiologie et Immunologie, Université de Montréal, Montréal, QC, Canada; Department of Biochemistry and Molecular Medicine, Université de Montréal, Montréal, QC, Canada; Department of Chemistry, School of Arts and Sciences, University of Pennsylvania, Philadelphia, PA, USA; Department of Biochemistry, Microbiology and Immunology, and Center for Infection, Immunity, and Inflammation, University of Ottawa, Ottawa, ON K1H 8M5, Canada

**Keywords:** VE607, COVID-19, SARS-CoV-2, SARS-CoV-1, spike glycoprotein, small molecule inhibitor, docking, single-molecule FRET

## Abstract

SARS-CoV-2 infection of host cells starts by binding of the Spike glycoprotein (S) to the ACE2 receptor. The S-ACE2 interaction is a potential target for therapies against COVID-19 as demonstrated by the development of immunotherapies blocking this interaction. Here, we present the commercially available VE607, comprised of three stereoisomers, that was originally described as an inhibitor of SARS-CoV-1. We show that VE607 specifically inhibits infection of SARS-CoV-1 and SARS-CoV-2 S-expressing pseudoviral particles as well as authentic SARS-CoV-2. VE607 stabilizes the receptor binding domain (RBD) in its “up” conformation. *In silico* docking and mutational analysis map the VE607 binding site at the RBD-ACE2 interface. The IC_50_ values are in the low micromolar range for pseudoparticles derived from SARS-CoV-2 Wuhan/D614G as well as from variants of concern (Alpha, Beta, Gamma, Delta and Omicron), suggesting that VE607 has potential for the development of drugs against SARS-CoV-2 infections.

## Introduction

The COVID-19 pandemic continues to cause widespread morbidity and mortality (Wu et al., 2020; Zhu et al., 2020a). This is largely due to insufficient vaccination levels as vaccines offer good protection against infection and severe disease (Ball, 2021). The currently used vaccines exploit modified versions of the Spike (S) glycoprotein that is exposed on the surface of viral particles (Krammer, 2020) and infected cells (Ding et al., 2022). S is processed by cellular proteases furin and TMPRSS2 on host cells. After binding to ACE2 via its receptor binding domain (RBD), S undergoes significant conformational changes that ultimately lead to fusion of the viral membrane with human cells. Fusion allows translocation of the RNA genome and associated replicase proteins into mammalian cells, leading to viral replication (Harrison et al., 2020; Hoffmann et al., 2020a; Hoffmann et al., 2020b; Yang and Rao, 2021). S is a trimeric glycoprotein that is present in multiple conformations that have been resolved primarily by cryo-electron microscopy (Cai et al., 2020; Lan et al., 2020; Shang et al., 2020; Wrapp et al., 2020; Yan et al., 2020). Its conformational dynamics can be monitored by single molecule FRET (Li et al., 2021b; Lu et al., 2020; Ullah et al., 2021; Yang et al., 2021). Vaccine-elicited antibodies act in several ways including neutralizing viral particles, but also through Fc-mediated effector functions (Tauzin et al., 2022; Tauzin et al., 2021). The selective pressure during the pandemic has led to a growing list of variants carrying mutations in the S-glycoprotein (Gong et al., 2021a; Li et al., 2021a; Mannar et al., 2021; Nabel et al., 2021; Prevost and Finzi, 2021; Yang and Rao, 2021) resulting in different degrees of resistance to previous infection and vaccine-elicited antibody neutralization.

Despite the efficacy of currently used vaccines and ongoing work to generate broadly protective pan-coronavirus vaccines (Cohen, 2021; Nabel et al., 2021; Rappazzo et al., 2021), there is an urgent need for efficient and specific treatments for infected patients. The viral replication machinery offers different possible drug targets (Yang and Rao, 2021) and several small molecule inhibitors targeting the SARS-CoV-2 protease (Dai et al., 2020; Zhang et al., 2020) or replicase (Kokic et al., 2021; Yin et al., 2021) have been published with some recently showing promise in clinical trials (Owen et al., 2021). In contrast, relatively little attention has been given to the S-ACE2 interaction as a potential target for small molecule inhibitors (Tong, 2009; Wang et al., 2021; Zhu et al., 2020b). Research on SARS-CoV-1 and Middle East respiratory syndrome (MERS) has inspired work on potential drug targets, and some previous studies explored the isolation of small molecule inhibitors against various potential targets. Some of these molecules were described as potential inhibitors of the SARS-CoV-1 RBD interaction with ACE2 (Adedeji et al., 2013; Kao et al., 2004), but the binding was not demonstrated directly and there was no biological follow-up work to characterize their mode of action.

Here we employed differential scanning fluorimetry (DSF) to identify the capacity of the small molecule inhibitor VE607 (Kao et al., 2004), composed of three stereoisomers, to bind the SARS-CoV-2 RBD. We found that this VE607 mixture of isomers (hereafter referred to as “VE607”) is capable of specific inhibition of infection of human cells with pseudoviral particles that express the SARS-CoV-1 or SARS-CoV-2 S-glycoproteins. VE607 was also able to inhibit the infection with authentic SARS-CoV-2 viruses. We found that VE607 inhibits the Spike by stabilizing the “up” conformation of the RBD. The mode of binding to RBD was elucidated by *in silico* docking experiments followed by validation of critical residues through mutagenesis and functional studies. Finally, VE607 remains potent against current variants of concern (VOC) of SARS-CoV-2 suggesting that it may be an interesting lead for the development of drugs for the prevention or treatment of COVID-19 infections.

## Results

### Differential scanning fluorimetry and docking suggest that VE607 may bind the RBD

We tested the ability of previously described SARS-CoV-1 inhibitors VE607 (Kao et al., 2004) and SSAA09E2 (Adedeji et al., 2013) to bind the SARS-CoV-2 RBD (Figure 1A). We used differential scanning fluorimetry (DSF) that, measures the effect of small molecules on the melting temperature of proteins (Mashalidis et al., 2013). Incubation with VE607 led to a significant decrease of the melting temperature (ΔT_m_, −2.3°C) while SSAA09E2 had a smaller, yet measurable effect (ΔT_m_, −0.7°C) (Figure 1B). Since this result suggested binding of VE607 to RBD, we next performed *in silico* docking against RBD using Glide (Schrödinger, 2020). We identified moderately favorable potential VE607 binding sites overlapping the ACE2 epitopes in both SARS-CoV-1 and SARS-CoV-2 RBDs (Figure 1C and D).

**Fig. 1.**
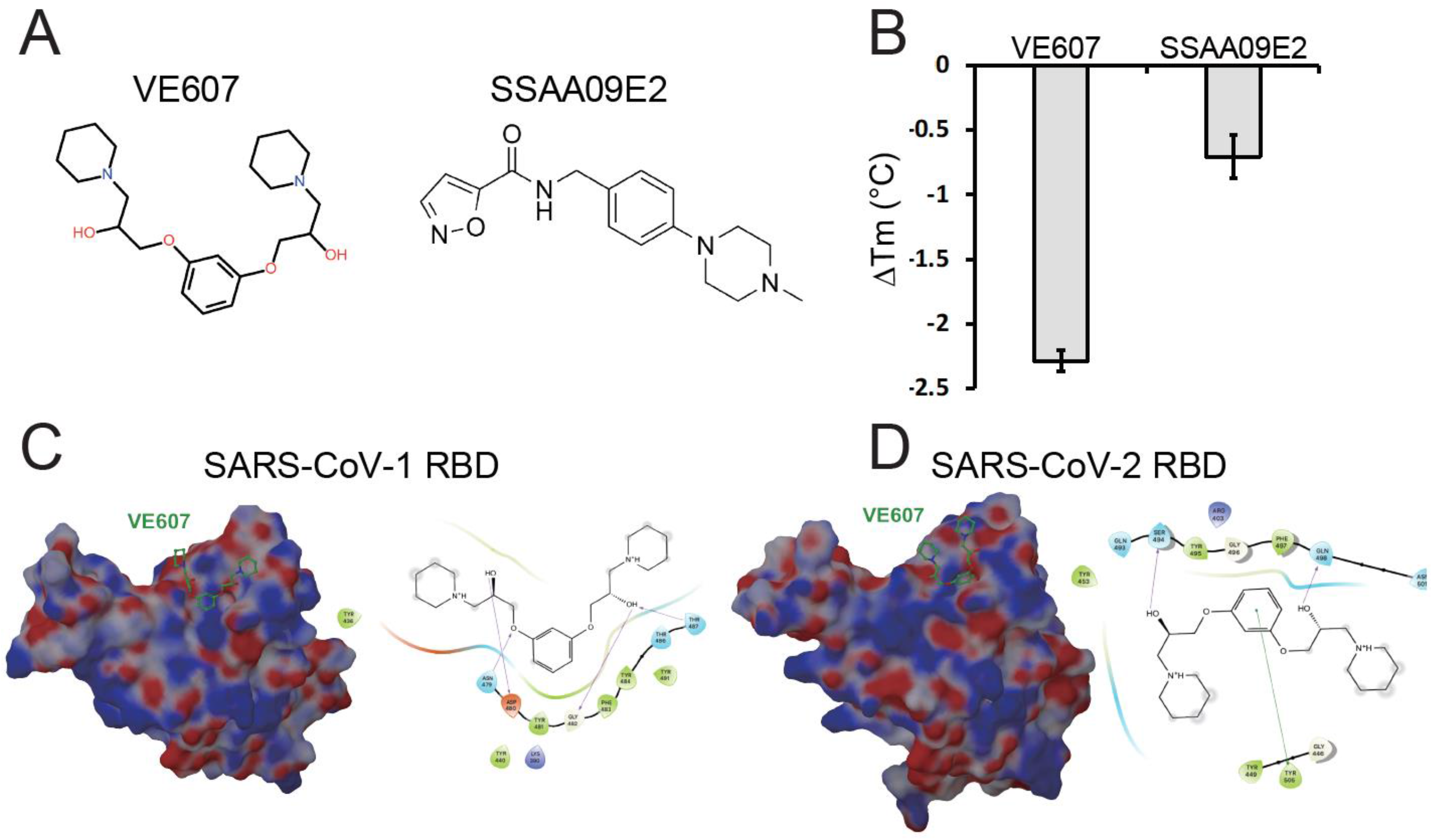
Potential interactions of SARS-CoV-1 inhibitors with the RBD. (A) Chemical structures of VE607 and SSAA09E2; (B) Differential scanning fluorimetry of the SARS-CoV-2 RBD in the presence of SARS-CoV-1 inhibitors, results from two experiments (eight replicates total) are shown; (C) Virtual docking of VE607 to SARS-CoV-1 and (D) SARS-CoV-2 RBD. Left panels, the electrostatic potential is displayed over molecular surface of the RBD and colored red and blue for negative and positive potential, respectively. Right panels, scheme showing a docking model of VE607 to the RBD. The presumable RBD contact residues are shown as spheres.

### VE607 inhibits infection of pseudoviral particles and authentic SARS-CoV-2

To assess the effects of VE607 and SSAA09E2 on infection we expressed the SARS-CoV-1 and SARS-CoV-2 S glycoproteins on the surface of pseudoviral particles carrying a luciferase reporter gene. Pseudoviral particles carrying the VSV-G glycoprotein served as control. Infection was measured using ACE2-expressing 293T (293T-ACE2) cells (Prevost et al., 2020) in the presence of increasing concentrations of VE607 and SSAA09E2. VE607 specifically inhibited pseudoviral particles bearing the SARS-CoV-1 Spike (IC_50_ = 1.47 μM, Figure 2A), in agreement with previous findings (Kao et al., 2004). Interestingly, VE607 also inhibited pseudoviral particles expressing the Spike from SARS-CoV-2 (IC_50_ = 3.06 μM, Figure 2A), albeit slightly less efficiently than for SARS-CoV-1. No inhibition was observed for pseudoviral particles bearing the VSV-G (IC_50_ > 100μM, Figure 2A). To ensure that the inhibitory capacity of VE607 against SARS-CoV-2 was not limited to pseudoviral particles, we evaluated whether the inhibitory capacity of VE607 was maintained against authentic viruses. As shown in Figure 2B, VE607 inhibited authentic SARS-CoV-2 D614G with an IC_50_ of 2.42 μM. No cell toxicity of VE607 or the three different enantiomers was observed with concentrations up to 100 μM on 293T ACE2 cells or Vero-E6 cells (Figure 2C). In contrast, SSAA09E2 at concentrations up to 100 μM did not inhibit infection of pseudoviral particles (data not shown) and we did not further pursue work with this small molecule.

**Fig. 2.**
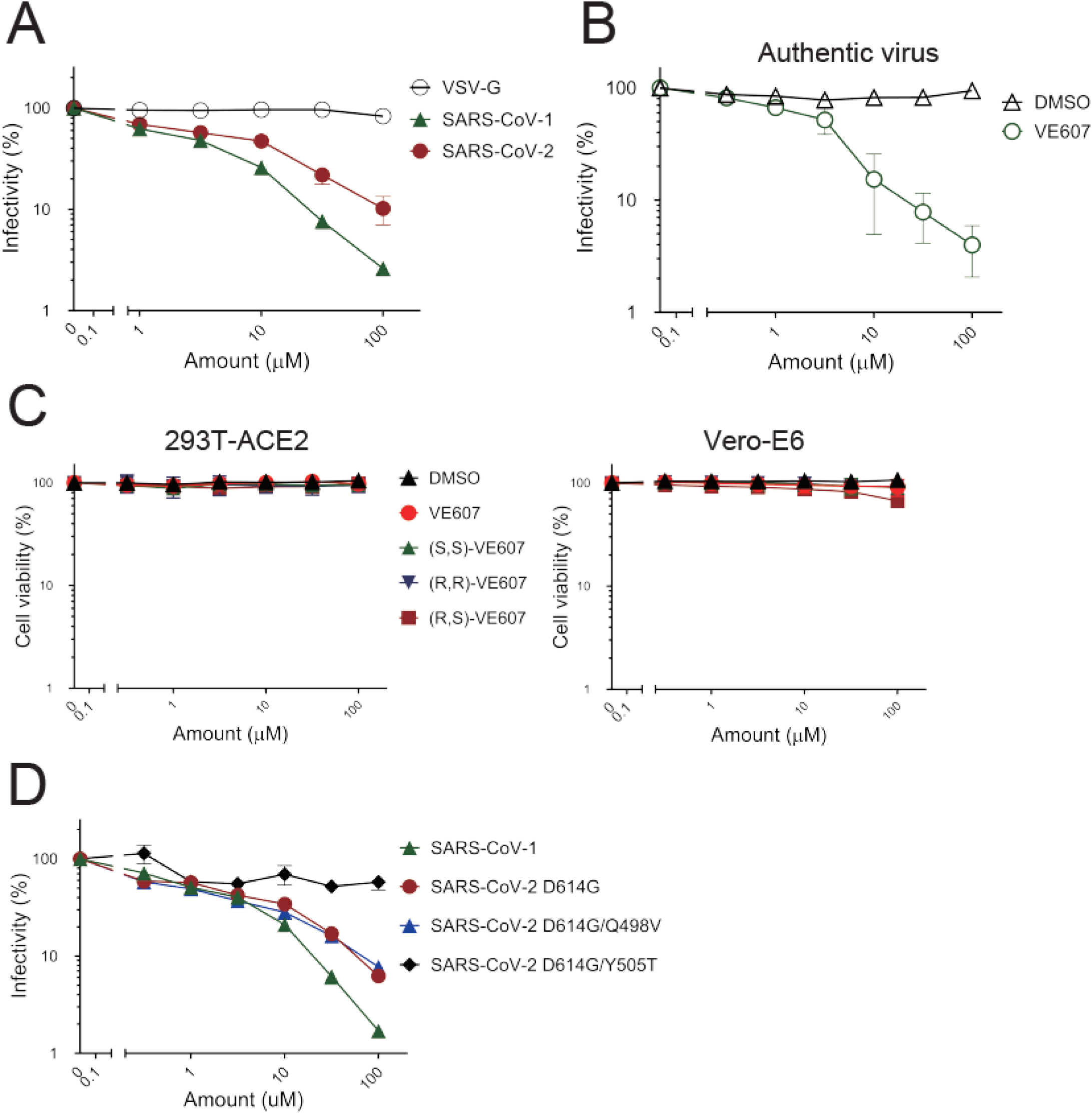
VE607 inhibits infection of SARS-CoV-1 and SARS-CoV-2 pseudoviral particles and of authentic SARS-CoV-2. (A) VE607 inhibition of SARS-CoV-1, SARS-CoV-2 or VSV-G (specificity control) pseudovirus; (B) VE607 inhibition of authentic live SARS-CoV-2 virus; (C) VE607 and the three different enantiomers are not toxic on 293T-ACE2 (left) or VERO-E6 (right) cells, as measured by CellTiter-Glo One Solution Assay for the quantitation of ATP presented in live cells. (D) Pseudovirus neutralization of SARS-CoV-2 S mutants predicted by our *in silico* analysis to modulate the inhibition by VE607. Data represents the average of at least four independent experiments ± SEM.

As stated above, commercially available VE607 is a mixture of three stereochemical isomers, comprised of the (S,S)-VE607, (R,R)-VE607, and the meso (R,S)-VE607. We observed no differences in the SARS-CoV-2 pseudoviral inhibition among these enantiomers obtained by synthesis, and the commercially available mixture of all three isomers (Figures S1).

Initial *in silico* docking identified RBD residues Y505 and Q498 as potential specific contact sites for VE607 (Figure 1D). We mutated these residues in the full-length SARS-CoV-2 D614G Spike and prepared pseudoviral particles to test whether they affect VE607 inhibition. While the Q498V mutation had only a minor effect (IC_50_ = 1.80 μM), the Y505T mutant was resistant to VE607 inhibition (IC_50_ > 40 μM, Figure 2D). These results are in agreement with the *in silico* analysis, where a strong π-π interaction between Y505’s aromatic side-chain and the central aromatic ring of VE607 is predicted. Alignment of sACE2 on the known ACE2 epitope of the VE607-bound model of RBD displayed significant steric clashes between ACE2 and VE607, suggesting some direct competition for the ACE2 epitope.

### VE607 stabilizes the “up” conformation of the S protein

We next assessed whether VE607 affects the RBD-ACE2 interaction. Briefly, we measured by flow cytometry the capacity of VE607 to compete with soluble ACE2 (sACE2) for interaction with the full SARS-CoV-2 Spike, expressed at the cell surface, as described previously (Anand et al., 2020). We observed no competition between VE607 (100 μM) and sACE2 (Figure 3A). Since the mode of action of some neutralizing antibodies such as CV3-1 involve S1 shedding (Li et al., 2022), we tested if VE607 acted in a similar manner. To this effect, Spike expressing 293T cells were radioactively labeled followed by the immunoprecipitation of cell lysates and supernatant, as described (Li et al., 2022). In contrast to CV3-1, VE607 decreased shedding, resulting in a significantly more stable trimer (Figure 3B).

**Fig. 3.**
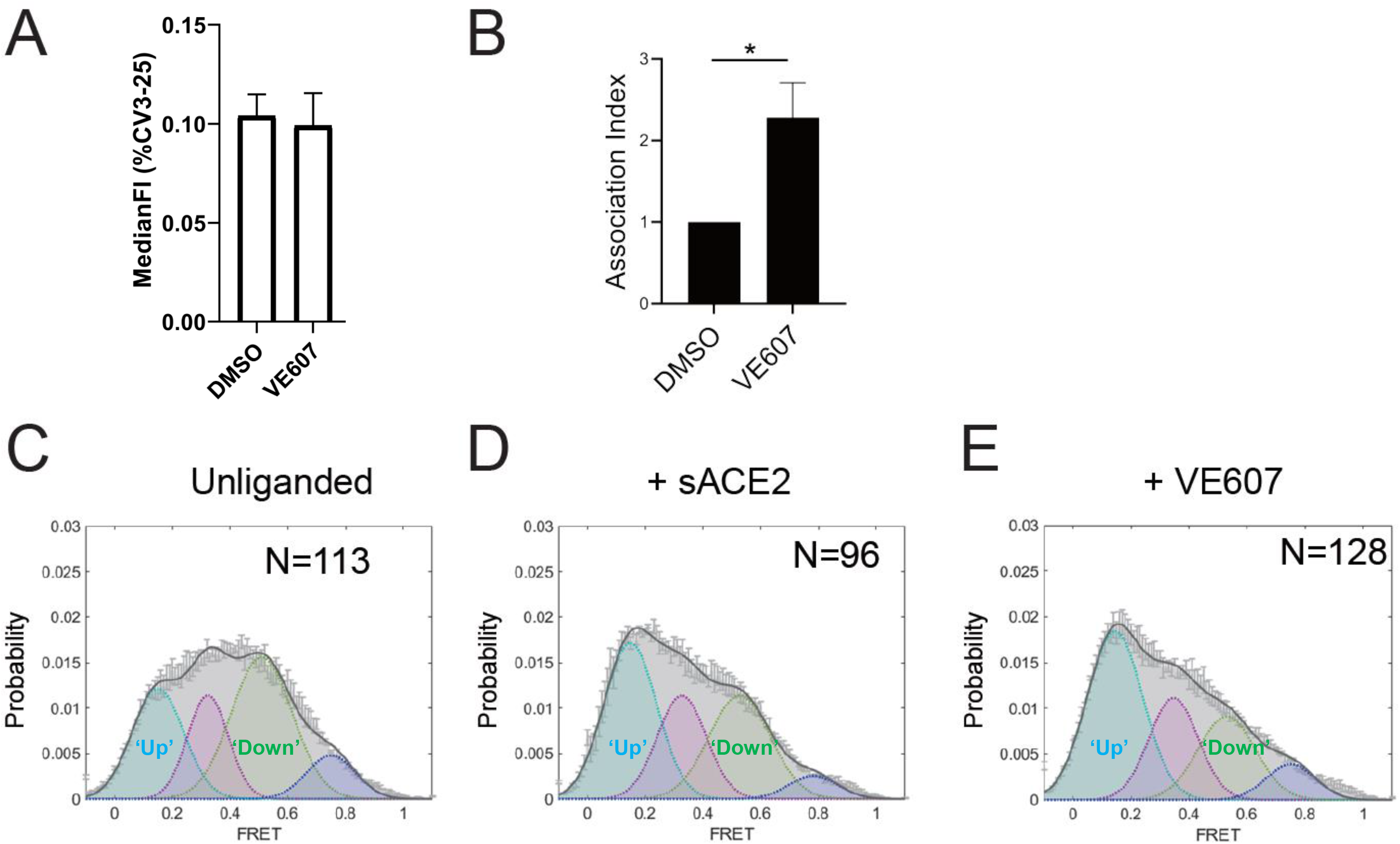
VE607 stabilizes SARS-CoV-2 S in the “up” conformation. (A) VE607 does not compete for sACE2 interaction, as measured by flow cytometry. The values represent the median fluorescence intensities (MFI) normalized to binding signals obtained with the conformationally independent CV3-25 Ab. Five experiments are represented as mean ± SEM and statistical significance was tested using unpaired t test. (B) SARS-CoV-2 Spike stability was measured by radioactive labelling of 293T Spike expressing cells followed by immunoprecipitation of cell lysates and supernatants. At least four experiments are represented as mean ± SEM and statistical significance was tested using unpaired t test, * *p* < 0.05. (C - E) Single molecule FRET analysis of SARS-CoV-2 S unliganded (C), in presence of sACE2 (D) or VE607 (E).

To evaluate whether the enhanced stability observed in presence of VE607 altered the conformational landscape of the Spike, we performed single-molecule FRET (smFRET) analysis using viral pseudoparticles carrying modified S glycoproteins labelled with FRET donor and acceptor dyes enabling us to distinguish the “up” and “down” conformations (Lu et al., 2020) (Li et al., 2022). In agreement with previous observations, the unliganded Spike predominantly sampled the “down” conformation (Figure 3C). As expected, the addition of sACE2 shifted the conformational landscape of the Spike to the “up” conformation reflecting the receptor bound state (Figure 3D). Interestingly, we observed that VE607 stabilized the “up” conformation mimicking sACE2 (Figure 3 E).

### VE607 inhibits infection of some SARS-CoV-2 variants of concern

SARS-CoV-2 is in constant evolution as VOCs keep emerging. VE607 was identified as an inhibitor of SARS-CoV-1, a related Beta-coronavirus, suggesting some inhibitory breadth. Therefore, we tested whether it inhibits pseudoviral particles bearing the Spike glycoproteins from the major VOCs (Alpha, Beta, Gamma, Delta and Omicron). In agreement with its broad SARS coronavirus activity, VE607 inhibited all VOCs with similar potency with IC_50_ values in the low micromolar range (Figure 4). These results demonstrate that the various amino acid changes in the S-glycoprotein of these variants do not impact the inhibitory potential of VE607 and show promise for the development a new generation of anti-SARS-CoV-2 small molecule inhibitors blocking viral entry.

**Fig. 4.**
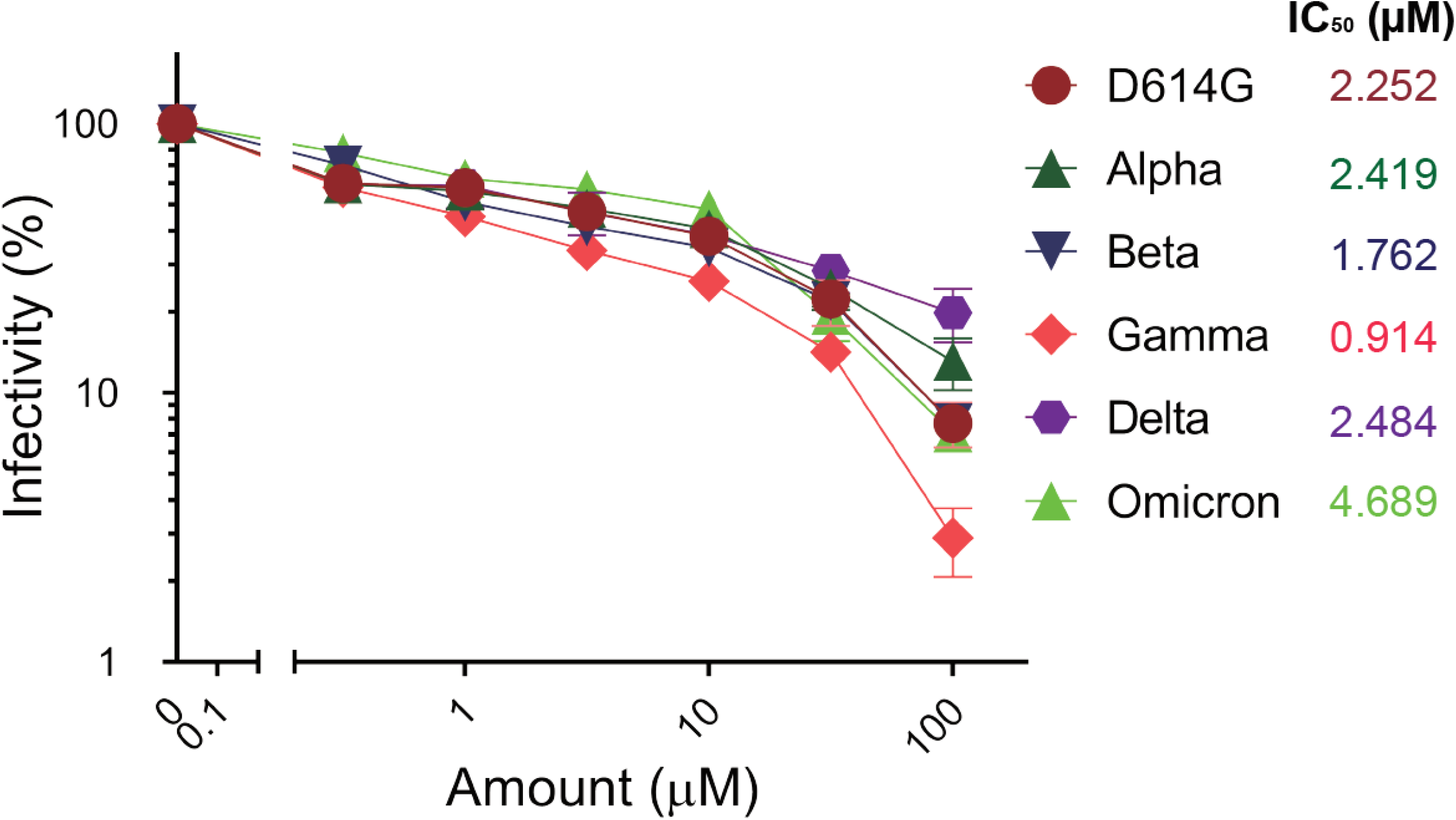
VE607 inhibits infection of SARS-CoV-2 variant Alpha, Beta, Gamma, Delta and Omicron pseudovirus particles. VE607 inhibits SARS-CoV-2 pseudoviral particles infection of 293T-hACE2 cells. IC_50_ values are shown next to the different VOCs Spikes. Data represents the average of at least four independent experiments ± SEM.

## Discussion

Here we present data suggesting that small molecule inhibitors of SARS-CoV-2 entry, such as VE607, have potential for the development of drugs for the prevention and/or treatment of COVID-19. VE607 was originally described as an inhibitor of the S glycoprotein-ACE2 mediated SARS-CoV-1 entry (Kao et al., 2004) and we confirmed these results using a pseudovirus infection assay. The inhibition of SARS-CoV-1 carrying pseudoviruses was stronger than that of SARS-CoV-2 pseudoviruses, but the IC_50_ values remained in the low μM range in both cases. *In silico* docking identified a potential binding site at the RBD-ACE2 interface in both cases (Lan et al., 2020; Shang et al., 2020). The results of mutational analysis are consistent with the predicted binding site suggesting a critical role of Y505 for the activity of VE607. Therefore, VE607 may inhibit viral entry by blocking ACE2-mediated Spike conformational changes required for fusion.

Single molecule FRET data indicate that VE607 alone stabilized the Spike in the preferred “up” conformation for sACE2 binding. However, in contrast to RBD-targeting antibodies such as CV3-1 (Li et al., 2022), VE607 did not induce shedding of S1. These results suggest a unique mechanistic basis for the inhibition of infection by VE607. We hypothesize that VE607 stabilizes one ACE2-bound conformation of Spike but is an allosteric inhibitor of downstream ACE2-triggered conformational changes required for fusion. The binding site predicted by *in silico* docking was confirmed by mutational analysis suggesting that VE607 binds at the S-ACE2 interface where it may block the protein-protein interaction required to activate the Spike for fusion. The emergence of variants over the course of the pandemic is a continuing concern and many of them carry mutations in the S glycoprotein including the RBD domain that contribute to increased infectivity and immune escape. To validate the binding site predicted by *in silico* docking we introduced the mutations Q498V and Y505T and whereas the first change had only a modest effect, the change Y505T led to an S glycoprotein that is resistant to VE607.

These results are consistent with the fact that pseudoviruses carrying S from the variants Alpha (B.1.1.7), Beta (B.1.351), Gamma (P.1), Delta (B.167.2) and Omicron (B.1.1.529) are inhibited by VE607 at low μM IC_50_ values, suggesting that common mutations in the vicinity of the binding site such as N501Y do not impact the inhibitory potential of VE607. Changes of Q498 are rarely observed in the Beta variant (less than 0.1% of all sequences), but the mutation Q498R is very frequent in Omicron (90% of all sequences) which is rapidly spreading and leading to the next phase of the pandemic (https://outbreak.info/compare-lineages). Interestingly, the Y505H mutation is also frequent in Omicron (90% of all sequences), but it is rarely present in other variants such as Alpha (less than 0.1% of all sequences). Nevertheless, the inhibitory effect of VE607 on pseudoviral particles carrying Spike from Omicron was comparable to other VOCs. It is presently unclear why the presence of Y505H does not affect the inhibitory activity of VE607 against Omicron when the Y505T change renders pseudoviral particles resistant to the molecule. It is possible that the role of Y505 in VE607 binding depends on the overall conformation of the Spike, which might differ in Omicron since it accumulated more than 30 mutations in the Spike. Altogether, our results constitute a proof of concept showing that small molecules targeting the SARS-CoV-2 Spike have potential for the development of drugs that may contribute to the fight against COVID-19.

## Acknowledgments

The authors thank the CRCHUM BSL3 and Flow Cytometry Platforms for technical assistance. The authors thank Hughes Charest and the LSPQ for the authentic SARS-CoV-2 virus. This work was supported by “Ministère de l’Économie et de l’Innovation du Québec, Programme de soutien aux organismes de recherche et d’innovation”, by the Fondation du CHUM, by a Canadian Institutes of Health Research (CIHR) foundation grant #352417 and by an Exceptional Fund COVID-19 from the Canada Foundation for Innovation (CFI) #41027 to A.F, and by R01 AI163395 from NIH NIAID to W.M. Work on the presented variants was also supported by the Sentinelle COVID Quebec network led by the LSPQ in collaboration with Fonds de Recherche du Québec Santé (FRQS) to A.F. The work was also supported by the UdeM Faculty of Medicine fund “Combattre la COVID-19: de la prévention au contrôle” to C.B. and A.F.; A.F. is the recipient of a Canada Research Chair on Retroviral Entry # RCHS0235 950-232424. G.B.B. is the recipient of a Fonds de Recherche Québec—Santé (FRQS) PhD fellowship. The funders had no role in study design, data collection and analysis, decision to publish, or preparation of the manuscript. The graphical abstract has been generated with BioRender website.

## Author Contributions

A.F. and C.B. designed the studies. S.D., S.Y.G., J.G., M.M., D.V., G.B.B., V.T.V., G.G., J.R., D.Y., A.B.S., M.P., M.C., C.A., W.M., A.F., and C.B., performed the experiments and interpreted the results. Y.C., A.B.S., M.P., and M.C. contributed unique reagents. A.F. S.D., and C.B. wrote the manuscript with inputs from others. Every author has read, edited and approved the final manuscript.

## DISCLAIMER

The views expressed in this manuscript are those of the authors and do not reflect the official policy or position of the Uniformed Services University, US Army, the Department of Defense, or the US Government.

## STAR Methods

### Lead Contact

Further information and requests for resources and reagents should be directed to and will be fulfilled by the Lead Contact.

### Materials Availability

All unique reagents generated during this study are available from the Lead contact with a completed Materials Transfer Agreement.

### Data and Code Availability

This study did not generate new code.

### Plasmids

The plasmids expressing the human coronavirus Spike of SARS-CoV-2 was kindly provided by Stefan Pöhlmann and was previously reported (Hoffmann et al., 2020b). The pNL4.3 R-E-Luc was obtained from NIH AIDS Reagent Program. The codon-optimized RBD sequence (encoding residues 319-541) fused to a C-terminal hexahistidine tag was cloned into the pcDNA3.1(+) expression vector and was reported elsewhere (Beaudoin-Bussieres et al., 2020). The plasmids encoding the SARS-CoV-2 variants Spikes D614G, Alpha (B.1.1.7), Beta (B.1.351), Gamma (P.1) were codon-optimized and synthesized by Genscript. Plasmid encoding the Delta (B.1.617.2) and Omicron (B.1.1.529) Spikes were generated by overlapping PCR using a codon-optimized wild-type SARS-CoV-2 Spike gene (GeneArt, ThermoFisher) that was synthesized (Biobasic) and cloned in pCAGGS as a template (Chatterjee et al., 2021; Gong et al., 2021b; Tauzin et al., 2022). The vesicular stomatitis virus G (VSV-G)-encoding plasmid (pSVCMV-IN-VSV-G) was previously described (Emi et al., 1991). Plasmids used to generate SARS-CoV-2 pseudoviral particles for smFRET analysis were described previously (Lu et. al., 2020).

### Cell lines

293T human embryonic kidney cells (obtained from ATCC) and Vero E6 cells (ATCC CRL-1586^™^) were maintained at 37°C under 5% CO2 in Dulbecco’s modified Eagle’s medium (DMEM) (Wisent) containing 5% fetal bovine serum (VWR), 100 UI/ml of penicillin and 100μg/ml of streptomycin (Wisent). The 293T-ACE2 cell line was previously reported (Prevost et al., 2020).

### Virus

Authentic SARS-CoV-2 was isolated, sequenced, and amplified from clinical samples obtained from patients infected with SARS-CoV-2 D614G by the Laboratoire de Santé Publique du Québec (LSPQ) and was previously described (Prevost et al., 2021). The virus was sequenced by MinION technology (Oxford Nanopore technologies, Oxford, UK). All work with the infectious SARS-CoV-2 authentic virus was performed in Biosafety Level 3 (BSL3) facilities at CRCHUM using appropriate positive-pressure air respirators and personal protective equipment.

### Methods detail

#### Purification of SARS-CoV-2 RBD

FreeStyle 293 F cells (Invitrogen) were grown in FreeStyle 293F medium (Invitrogen) to a density of 1 × 10^6^ cells/mL at 37°C with 8% CO_2_ with regular agitation (150 rpm). Cells were transfected with a plasmid coding for SARS-CoV-2 S RBD using ExpiFectamine 293 transfection reagent, as directed by the manufacturer (Invitrogen). One week later, cells were pelleted and discarded. Supernatants were filtered using a 0.22 μm filter (Thermo Fisher Scientific). The recombinant RBD proteins were purified by nickel affinity columns, as directed by the manufacturer (Invitrogen). The RBD preparations were dialyzed against phosphate-buffered saline (PBS) and stored in aliquots at −80°C until further use. To assess purity, recombinant proteins were loaded on SDS-PAGE gels and stained with Coomassie Blue.

#### Differential scanning fluorimetry

DSF experiments were essentially performed as described previously (Sharifahmadian et al., 2017). DSF was conducted using 5 μM of purified RBD, 10x concentration of SYPRO Orange (from 5000x stock solution, ThermoFisher) in 50 mM HEPES, 100 mM NaCl, pH 7.5 and 5% final concentration of DMSO. The small molecules were added to final concentrations of 5 mM. SYPRO Orange fluorescence was monitored over 20–95 °C with a LightCycler^®^ 480 instrument (Roche, USA). The LightCycler^®^ 480 Software (Roche) was used to calculate the first derivate of the resulting melting curve, with the steepest point of the slope being the Tm.

#### Molecular modeling

System preparation, modeling, and docking calculation were performed using the Schrödinger Suite package (Schrödinger, 2020), using default parameters unless otherwise noted. The target structures were taken from SARS-CoV-1 RBD (PDB ID: 6waq) and SARS-CoV-2 RBD (PDB ID: 6w41) prepared using the Protein Preparation Wizard (Sastry et al., 2013). To prepare the structures, force field atom types and bond orders were assigned, missing atoms and side-chains were added, protonation states of ionizable amino acid side-chains were determined using PROPKA (Olsson et al., 2011), water orientations were sampled, and hydrogen bond networks were subsequently optimized by flipping Asn/Gln/His residues and sampling hydroxyl/thiol hydrogen. Constrained energy minimization was then performed using the imperf module from impact (Schrödinger, 2020) to generate the structure to be used in the subsequent modeling calculations. Potential binding sites were explored and characterized using the SiteMap tool (Halgren, 2007; Halgren, 2009). VE607 compound was structurally preprocessed using LigPrep (Schrödinger, 2020) to generate multiple states for stereoisomers, tautomers, ring conformations, and protonation states at a selected pH range. Then, energy minimization was performed with the OPLS3e force field (Roos et al., 2019). The prepared molecular structures were docked into the putative binding sites using Glide (Friesner et al., 2004; Halgren et al., 2004) with the standard precision (SP) scoring function to evaluate enrichment of the calculated docked models.

#### Neutralization assay using pseudoviral particles

Target cells were infected with single-round luciferase-expressing lentiviral particles as described previously (Prevost et al., 2020). Briefly, 293T cells were transfected by the calcium phosphate method with the lentiviral vector pNL4.3 R-E-Luc (NIH AIDS Reagent Program) and a plasmid encoding for SARS-CoV-2 Spike at a ratio of 5:4. Two days post-transfection, cell supernatants were harvested and stored at –80°C until use. 293T-ACE2 target cells were seeded at a density of 1×10^4^ cells/well in 96-well luminometer-compatible tissue culture plates (Perkin Elmer) 24h before infection. Recombinant viruses in a final volume of 100μl were incubated with the indicated concentrations of small molecules (VE607 or SSAA009E2) up to concentrations of 100 μM for 1h at 37°C and were then added to the target cells followed by incubation for 48h at 37°C; cells were lysed by the addition of 30μl of passive lysis buffer (Promega) followed by one freeze-thaw cycle. An LB941 TriStar luminometer (Berthold Technologies) was used to measure the luciferase activity of each well after the addition of 100μl of luciferin buffer (15mM MgSO_4_, 15mM KPO_4_ [pH 7.8], 1mM ATP, and 1mM dithiothreitol) and 50μl of 1mM d-luciferin potassium salt (Prolume). The neutralization half-maximal inhibitory dilution (ID_50_) represents the sera dilution to inhibit 50% of the infection of 293T-ACE2 cells by recombinant viruses.

#### Cell surface staining and flow cytometry analysis

Using the standard calcium phosphate method, 10 μg of Spike expressor and 2.5 μg of a green fluorescent protein (GFP) expressor (pIRES-GFP) were transfected into 2 × 10^6^ 293T cells. 48h post-transfection Spike-expressing cells were incubated with 100 μM of VE607 or equivalent volume of vehicle (DMSO) and incubated for 30 min at room temperature. CV3-25 (5 μg/ml) or sACE2 (100 μg/ml) was added to the cells and incubated for 45min at 37°C and sACE2 binding was detected using a polyclonal Goat anti-human ACE2 (RND Systems) at 1/100 dilution at room temperature for 30min. AlexaFluor-647-conjugated goat anti-human IgG (H+L) Ab (Invitrogen) and AlexaFluor-conjugated donkey anti-goat IgG (H+L) Ab (Invitrogen) was used as secondary antibodies. The percentage of transfected cells (GFP+ cells) was determined by gating the living cell population based on viability dye staining (Aqua Vivid, Invitrogen). sACE2 binding levels were normalized to signals obtained with the conformationally independent anti-S2 CV3-25 mAb (Li et al., 2022; Prevost et al., 2021; Tauzin et al., 2022). Samples were acquired on a LSRII cytometer (BD Biosciences, Mississauga, ON, Canada) and data analysis was performed using FlowJo vX.0.7 (Tree Star, Ashland, OR, USA).

#### Radioactive labeling and immunoprecipitation

For pulse-labeling experiments, 5 × 10^5^ 293T cells were transfected by the calcium phosphate method with SARS-CoV-2 D614G Spike expressor. One day after transfection, cells were metabolically labeled for 16-24 h with 100 μCi/ml [^35^S]methionine-cysteine ([^35^S] protein labeling mix; Perkin-Elmer) in Dulbecco’s modified Eagle’s medium lacking methionine and cysteine and supplemented with 10% of dialyzed fetal bovine serum and 1% GlutaMAX^™^. Simultaneously, cells were treated with or without 100 μM VE607. Cells were subsequently lysed in radio immunoprecipitation assay (RIPA) buffer (140 mM NaCl, 8 mM Na_2_HPO_4_, 2 mM NaH_2_PO_4_, 1% IGEPAL^®^ CA-630, 0.05% sodium dodecyl sulfate [SDS], 1.2 mM sodium deoxycholate [DOC]). Precipitation of radiolabeled SARS-CoV-2 D614G Spike glycoprotein from cell lysates or supernatant was performed with CV3-25 and polyclonal rabbit antiserum raised against SARS-CoV-2 RBD protein, for 1 h at 4°C in the presence of 45 μL of 10% protein A-Sepharose beads (GE Healthcare). Samples were washed twice with RIPA buffer and then boiled 5 min in Laemmli buffer with β-mercaptoethanol before being separated by SDS-PAGE. After migration, gels were dried with a Model 583 gel dryer (Bio-Rad) and exposed to a storage phosphor screen. Densitometry data were acquired with a Typhoon Trio Variable Mode Imager (Amersham Biosciences) in storage phosphor acquisition mode and analyzed using ImageQuant 5.2 (Molecular Dynamics). Association index was determined by precipitation of radiolabeled cell lysates and supernatants with CV3-25 and polyclonal rabbit antiserum raised against SARS-CoV-2 RBD protein. The association index is a measure of the ability of the VE607 treated S1 subunit to remain associated with the trimeric spike (S) protein on the expressing cell relative to that of the mock-treated S1 and was calculated with the following formula: association index = ([cell S1]_treated_/[supernatant S1]_treated_)/([cell S1]_mock-treated_/[supernatant S1]_mock-treated_).

#### Microneutralization with authentic virus

One day prior to infection, 2×10^4^ Vero E6 cells were seeded per well in the 96-well flat bottom plate and incubated overnight to permit Vero E6 cell adherence. Compounds dilutions ranged from 0, 0.316, 1, 3.16, 10, 31.6 and 100 μM were performed in a separate 96 well culture plate using DMEM supplemented with penicillin (100 U/mL), streptomycin (100 μg/mL), HEPES, 0.12% sodium bicarbonate, 2% FBS and 0.24% BSA. 10^4^ TCID_50_/mL of SARS-CoV-2 virus was prepared in DMEM + 2% FBS and combined with an equivalent volume of diluted compounds for one hour. After this incubation, all media was removed from the 96 well plate seeded with Vero E6 cells and virus: compounds mixture was added to each respective well at a volume corresponding to 600 TCID_50_ per well and incubated for one hour further at 37°C. Both virus only and media only (MEM + 2% FBS) conditions were included in this assay. All virus-compounds supernatant was removed from wells without disrupting the Vero E6 monolayer. Each diluted compound (100 μL) was added to its respective Vero E6-seeded well in addition to an equivalent volume of MEM + 2% FBS and was then incubated for 48 hours. Media was then discarded and replaced with 10% formaldehyde for 24 hours to cross-link Vero E6 monolayer. Formaldehyde was removed from wells and subsequently washed with PBS. Cell monolayers were permeabilized for 15 minutes at room temperature with PBS + 0.1% Triton X-100, washed with PBS and then incubated for one hour at room temperature with PBS + 3% non-fat milk. An anti-mouse SARS-CoV-2 nucleocapsid protein (Clone 1C7, Bioss Antibodies) primary antibody solution was prepared at 1 μg/mL in PBS + 1% non-fat milk and added to all wells for one hour at room temperature. Following extensive washing (3×) with PBS, an anti-mouse IgG HRP secondary antibody solution was formulated in PBS + 1% non-fat milk. One hour post-room temperature incubation, wells were washed with 3× PBS, substrate (ECL) was added and an LB941 TriStar luminometer (Berthold Technologies) was used to measure the signal each well.

#### Cell viability test

To measure the cytotoxicity of VE607 and its stereoisomers on 293T-ACE2 or Vero-E6 cells, a cell viability assay using CellTiter-Glo^®^ One Solution Assay (Promega) was performed. Briefly, 293T-ACE2 or Vero-E6 cells were seeded at a density of 1×10^4^ cells/well in 96-well luminometer-compatible tissue culture plates (Perkin Elmer); After 24h, indicated concentrations of VE607, (S,S)-VE607, (R,R)-VE607 or (R,S)-VE607 up to concentrations of 100 μM were added to the cells followed by incubation for 48h at 37°C, same volume of its vehicle, DMSO, was added as control. Then a volume of CellTiter-Glo^®^ One Solution equal to the volume of cell culture medium present in each well was added, followed by 2 min mixing on shaker and 10 min incubation at room temperature. An LB941 TriStar luminometer (Berthold Technologies) was used to measure the luciferase activity of each well.

#### Chemical synthesis of the three enantiomers of VE607

All reactions were conducted in oven-dried glassware under an inert atmosphere of nitrogen, unless otherwise stated. All solvents were reagent or high-performance liquid chromatography (HPLC) grade. Anhydrous THF was obtained from the Pure SolveTM PS-400 system under an argon atmosphere. All reagents were purchased from commercially available sources and used as received. Reactions were magnetically stirred under a nitrogen atmosphere, unless otherwise noted and were monitored by thin layer chromatography (TLC) was performed on pre-coated silica gel 60 F-254 plates (40-55 micron, 230-400 mesh) and visualized by UV light or staining with KMnO4 and heating. Yields refer to chromatographically and spectroscopically pure compounds. Optical rotations were measured on a JASCO P-200 polarimeter. Proton (^1^H) and carbon (13C) NMR spectra were recorded on a Bruker Avance III 500-MHz spectrometer. Chemical shifts (δ) are reported in parts per million (ppm) relative to chloroform (δ 7.26) or methanol (δ 3.31) for ^1^H NMR, and chloroform (δ 77.2) or methanol (δ 49.0). High resolution mass spectra (HRMS) were recorded at the University of Pennsylvania Mass Spectroscopy Service Center on either a VG Micromass 70/70H or VG ZAB-E spectrometer. Lyophilization was performed in a Labconco FreeZone 12 Plus lyophilizer (0.148 mbar). The purity of new compounds were judged by NMR and LCMS (>95%).

#### smFRET analysis

Pseudoviral particles bearing labeled CoV2_WH01_ S protein were prepared, imaged, and analyzed as described previously (Li et al., 2021c; Lu et al., 2020). Samples were pre-incubated with sACE2 (200 μg/ml) or VE607 (100 μM) for 90 minutes at room temperature prior to imaging.

#### Quantification and statistical analysis

Statistics were analyzed using GraphPad Prism version 8.0.2 (GraphPad, San Diego, CA, (USA). Every data set was tested for statistical normality and this information was used to apply the appropriate (parametric or nonparametric) statistical test. P values <0.05 were considered significant; significance values are indicated as * p<0.05; ** p<0.01; *** p<0.001; **** p<0.0001.

---

**Supplemental Figure 1.**
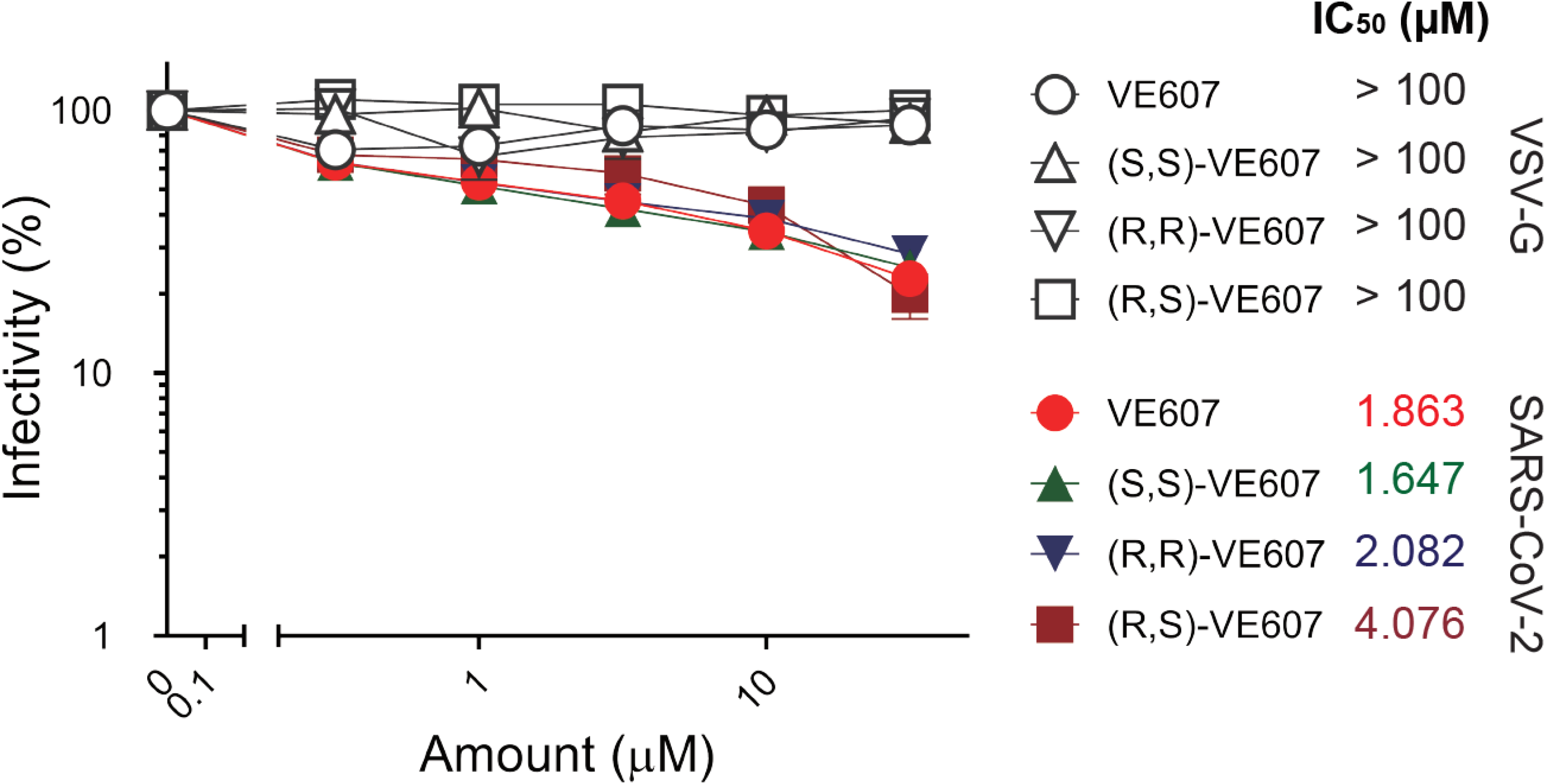
Stereochemical isomers of VE607 have the same capacity as commercially available VE607 to neutralize SARS-CoV-2 pseudovirus particles. All three stereochemical isomers of VE607, (S,S)-VE607, (R,R)-VE607 and (R,S)-VE607 were tested for their inhibition of VSV-G pseudovirus or SARS-CoV-2 D614G pseudovirus. IC50 values are shown next to the different VOC Spikes. Data represents the average of at least four independent experiments ± SEM.

